# Modeling a Continuum of Drug-induced Persistence during Targeted Cancer Therapy

**DOI:** 10.1101/2025.01.21.634165

**Authors:** Ji Tae Park, Herbert Levine

**Affiliations:** Department of Physics, Northeastern University, Boston, MA 02115, USA; Center for Theoretical Biological Physics, Northeastern University, Boston, MA 02115, USA; Department of Bioengineering, Northeastern University, Boston, MA 02115, USA

## Abstract

We introduce a general phenomenological framework for understanding how phenotypic plasticity gives rise to drug persisters. These persisters, often quiescent but sometimes which again return to cycling, survive in the presence of targeted therapy and eventually can lead to mutants with true resistance. Our framework builds on recent experimental observations regarding variations between and among single-cell clones and the possible role of the drug itself in enhancing the survival strategy. Predictions of our approach include the existence of an optimum drug concentration as well as an optimum drug holiday schedule to minimize the persistence-based threat.

## 1 Introduction

Despite ongoing advancements, discoveries, and drug developments in oncology, many patient outcomes remain suboptimal, with refractory and recurrent diseases continuing to result in substantial morbidity and mortality worldwide [1–3]. Although a plethora of mechanisms of resistance to specific therapies have been studied and identified [4], our understanding of the fundamental principles that allow cancer cells to adapt to stressful environments, such as drug exposure, remains incomplete. Traditionally, resistance to anticancer agents has been attributed to stochastic genetic mutations that alter drug targets, create specific and/or non-specific efflux pumps, or induce other advantageous changes in biochemical pathways [5]. However, recent research has highlighted a different phenomenon known as persistence – distinct from resistance – which is believed to be a reversible, drug-tolerant state achieved through non-mutation-based adaptation, and typically associated with a low proliferation rate [6].

Persistence, first discovered and extensively studied in bacterial systems [7, 8], has only recently garnered attention among cancer biologists. Unlike resistance, whether pre-existent or later acquired by stochastic genetic mutations, persistence is achieved exclusively through shifts in phenotypic states [9]. Indeed, the phenotypic plasticity of cancer cells have long been considered one of the key driving factors behind acquired drug tolerance [10], minimal residual disease (MRD) and dormancy [11], the epithelial-mesenchymal transition (EMT) [12], and cancer stem cell (CSC) formation [13]. A growing body of experimental evidence suggests that upon exposure to a drug, a subpopulation of cancer cells can transiently enter a drug-tolerant state without any alterations in genotype and can regain sensitivity once the drug is removed [14, 15]. Furthermore, although these drug-tolerant persisters (DTP) are primarily found to be quiescent, a small fraction of them can re-enter the proliferation cycle, thereby forming “drug-tolerant expanding persisters” (DTEP) also known as “cycling persisters” [14, 16, 17].

There is also increasing evidence linking persistence to adverse treatment outcomes. Hata et al. [18] reported that a subset of drug-tolerant PC9 human non-small cell lung carcinoma (NSCLC) cells subsequently underwent a T790M mutation, conferring resistance. This occurred after prolonged exposure to gefitinib, an anti-EGFR tyrosine kinase inhibitor (TKI). Similarly, Ramirez et al. [19] demonstrated that extended treatment with erlotinib, another anti-EGFR TKI, can eventually lead to the *de novo* acquisition of resistance by persisting PC9 cells, through diverse mechanisms. Furthermore, Russo et al. [20] suggested that persistent cells exhibit stress-induced mutagenesis, increasing their likelihood of acquiring resistance compared to drug-sensitive cells, based on their experiments exposing colorectal cancer cells to either dabrafenib, an anti-BRAF TKI, and cetuximab, an anti-EGFR monoclonal antibody, or to cetuximab alone to induce persistence. They then employed the Luria-Delbrück fluctuation assay to quantify rates of resistance development [20]. Notably, their mathematical model, which represents sensitive and persistent cells as two discrete sets of population with drug concentration-dependent transition rates between them, most accurately fit the group’s empirical data only after assuming that persistence is largely, if not entirely, drug-induced [20]. This also agrees with their observed Poisson abundance distribution of persisters [20]. This finding notably contrasts with bacterial systems, where a small population of (potentially) persistent cells is thought to be typically maintained through bet-hedging strategies [21]. Collectively, many experimental observations underscore cancer cells’ phenotypic plasticity that enables extensive adaptations to various environmental stressors, such as cytotoxic drugs.

In this study, we develop a mathematical framework to describe the dynamic process by which cancer cells enter a persistent state and predict their subsequent fates under treatment. Our approach conceptualizes epigenetic and phenotypic states as one-dimensional continua, and it models the population probability distribution of these cells as temporally evolving on this space due to both random (diffusive) and adaptive (drug-induced) processes in addition to growth and death terms. All of these effects depend on drug concentration, which itself can be a function of time. As we will see, extending the concept of persistence from a discrete (sensitive versus persistent) [22–24] to a continuous phenotype enables contact with new experimental protocols.

Our approach non-trivially extends previous work by Kessler and Levine [25] (see also [16] and [26]) who employed a phenomenological model to characterize persistence as observed in Berger et al.’s experiments on PC9 cells treated with gefitinib [27]. In their experiments, Berger et al. measured the chance-to-persist (CTP), defined as the survival fraction on day 7 of the treatment, for each single-cell clone derived from the cell line. This measurement clearly demonstrated the continuous nature of persistence. Kessler and Levine [25] developed a partial differential equation (PDE) computational model that describes the temporal progression in a one-dimensional epigenetic space regulating a clone’s CTP. Their modeling results align well with the experimental data, but there were several important deficiencies. First, the model lacked concentration-dependent mechanisms by which drug presence specifically induces persistence. Furthermore, the model contains no mechanism allowing for the re-emergence of proliferation in the population via cycling persisters.

To address these limitations, we incorporate new experimentally motivated processes into the PDE framework. Our approach predicts a critical CTP value for a given drug concentration that can serve as a reference for forecasting a persisting clone’s eventual fate – gradual extinction or cycling persistence – and allows us to explore different treatment schemes. In particular, we evaluate the potential benefits of intermittent drug holidays. We emphasize that our framework is not intended to be a set of specific mathematical models designed to fit specific experimental data points. Instead, it provides a broad theoretical framework that enables us to investigate more general concepts and principles.

## 2 Results

### 2.1 The Model

We assume that within a tissue or population of cancer cells sharing the same genotype, each single-cell clone exhibits a distinct probability of persisting under treatment. To quantify this, we adopt an approach inspired by Berger et al. [27] and Kessler and Levine [25], operationally defining CTP, denoted as *x* ∈ (0, 1), for a given clone as the surviving fraction of the population after continuous drug treatment at a reference constant concentration *c*(*t*) = *c*_*p*_ for a duration *τ*. In our context, we set *c*_*p*_ = 1 (in arbitrary units) and *τ* = 7 days. We interpret *x* as specifying a long-lived epigenetic state that determines a clone’s corresponding CTP [14, 28, 29]. This state can dynamically change, albeit a slow rate, given by the parameter *µ*, allowing cells to explore different epigenetic states (i.e. different values of CTP). We also assume that increasing the CTP (*x*) incurs a fitness cost, which manifests as a reduction in the clone’s proliferation rate in the absence of any drug.

Sharing the same epigenetic state does not imply complete phenotypic homogeneity among cells. In fact, phenotypic heterogeneity, coupled with plasticity, has been shown to enhance the progression of cancer cells, driving non-genetically acquired tolerance and eventual resistance to therapies [30]. To account for this additional variability, we introduce a second continuous variable, *s* ∈ ℝ, representing phenotypic states associated with survival under stress. Cells with higher *s* values display greater survival than those with lower *s* values. Each epigenetic state *x* determines an initial (i.e. before treatment) clonal population distribution in the phenotypic space *s* – a Gaussian distribution around the mean *s*_0_(*x*) (Figure 1(a)). Large *x* translates to an initial distribution more skewed toward better survival 1(a). Our model allows the treatment to induce phenotypic adaptation and dynamically alter this distribution. Effectively, tumor cells can be viewed as diffusing particles on the *x*-*s* plane under a drug-dependent potential and birth and death events.

**Figure 1:**
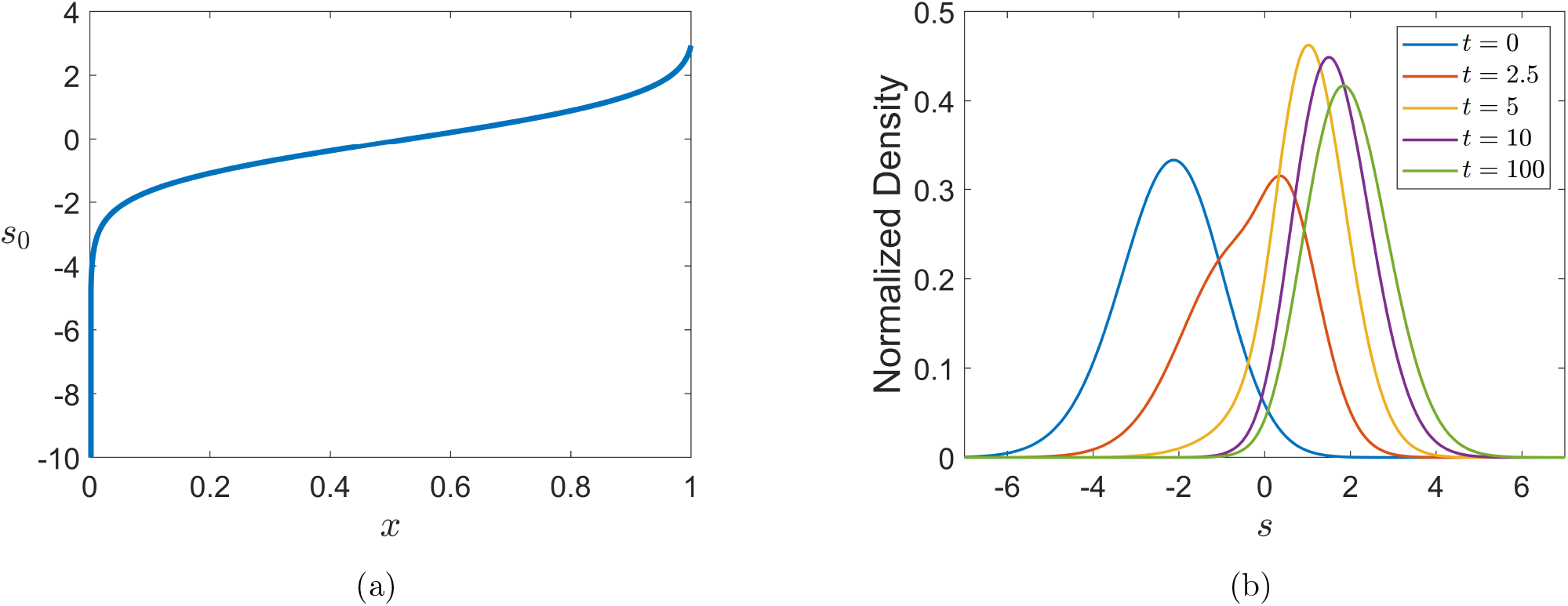
(a) The plot of *s*_0_(*x*) as a function of *x*. The initial distribution in *s* for a clone with a fixed *x* is precisely determined by *s*_0_(*x*). (b) Plots of the normalized distributions in *s* subjected to both advection (*v*) and selection (*r* − *k*). The parameters used for these plots can be found in Table S1 (Supporting Information).

Our model consists of diffusive terms (in both *x* and *s*), an advective term *v*(*s, c*(*t*)), a killing term *k*(*s, c*(*t*)), and a proliferative term *r*(*x, s, c*(*t*)) (see Materials and Methods). Partially inspired by Clairambault et al. [31], the advection captures the drug-induced states within the phenotypic space. This directed stress adaptation is corroborated by experimentally observed sequential cell state transitions from sensitive to quiescent persistent, and then to cycling persistent states [14, 17]. Simultaneously, this term highlights the paradoxical effect of treatment; while drugs induce cytotoxicity, they also promote phenotypic adaptations that enable persistence. In our model, we assume that *v*(*s, c*(*t*)) has an asymptotic upper limit of *v*_0_, which saturates sigmoidally toward *v*_*max*_ with increasing drug concentration, *c*(*t*), with cooperativity of *h*. This formulation prevents the advection rate from becoming arbitrarily and unrealistically large at high drug doses. Additionally, *v*(*s, c*(*t*)) is a decreasing function of *s*, reflecting the assumption that cells have a maximal amount of adaptability, perhaps occasioned by the fact that cells with sufficient *s* do not require further advection to gain a survival advantage in typical challenging environments. Alternatively, this may indicate an inherent limit to the degree of adaptation due to intrinsic physiological factors.

Analogous to the advection term, the killing rate (*k*(*s, c*(*t*))) as a function of *s* is bounded from above by *k*_0_, which itself asymptotically approaches *k*_*max*_ as *c*(*t*) increases. We constructed the form of *k*_0_(*t*) to resemble a dose-response curve with *c*_50_ as the drug concentration at which 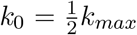. The killing rate diminishes at larger *s* values, approaching zero as cells become more drug-tolerant.

Lastly, the proliferation term (*r*(*x, s, c*(*t*))) is influenced by both epigenetic (*x*) and phenotypic (*s*) states. The function *r*(*x, s, c*(*t*)) is a slowly increasing logistic function in *s*, bounded from above by *r*_0_. It is important to note that quiescence is not a consequence of the balance between birth and death terms (*r* − *k* ≈ 0), but instead it is a state marked by near, if not complete, cell-cycle arrest [32]. The proliferation is also dependent on *x*, with clones having a baseline growth rate of *ξ* while incurring a cost for obtaining higher CTP. Moreover, the total population globally attempts to reach its carrying capacity, *N*_*ss*_, the initial steady-state population before treatment. Notably, *r*(*x, s, c*(*t*)) becomes primarily determined by *x* (*r* ≈ *R*_*x*_(*x, t*)) in the (near) absence of stress (drug), while it is predominantly affected by *s* (*r* ≈ *R*_*s*_(*s, t*)) in the presence of stress (drug).

Although the population distribution, *P* (*x, s, t*), across epigenetic phenotypic spaces can, in principle, evolve in both *x* and *s*, we assume that changes in *s* occur on a much shorter timescale than changes in *x*. The assumption is justified as phenotypic transitions typically occur rapidly, whereas epigenetic changes are inherently slower. To reflect this, we impose the condition that diffusion in *s* is much faster than in *x*. This ensures that the *s*-dynamics dominate in the presence of a drug (*c*(*t*) *>* 0) while the *x*-dynamics play a more prominent role in the absence of a drug (*c*(*t*) = 0).

A more detailed description of the model can be found in Materials and Methods.

### 2.2 Continuous Treatment

For a fixed set of parameters (see Table S1 in Supporting Information), we first performed a 100-day continuous treatment simulation with *c*(*t*) = 1 and generated survival curves (*A*(*x, t*)) for given values of *x* (Figure 2(a)). We then also generated survival curves for the entire population, *N* (*t*), at different constant drug concentrations ranging from 10^*−*4^ to 10^1^ (Figure 2(b)).

**Figure 2:**
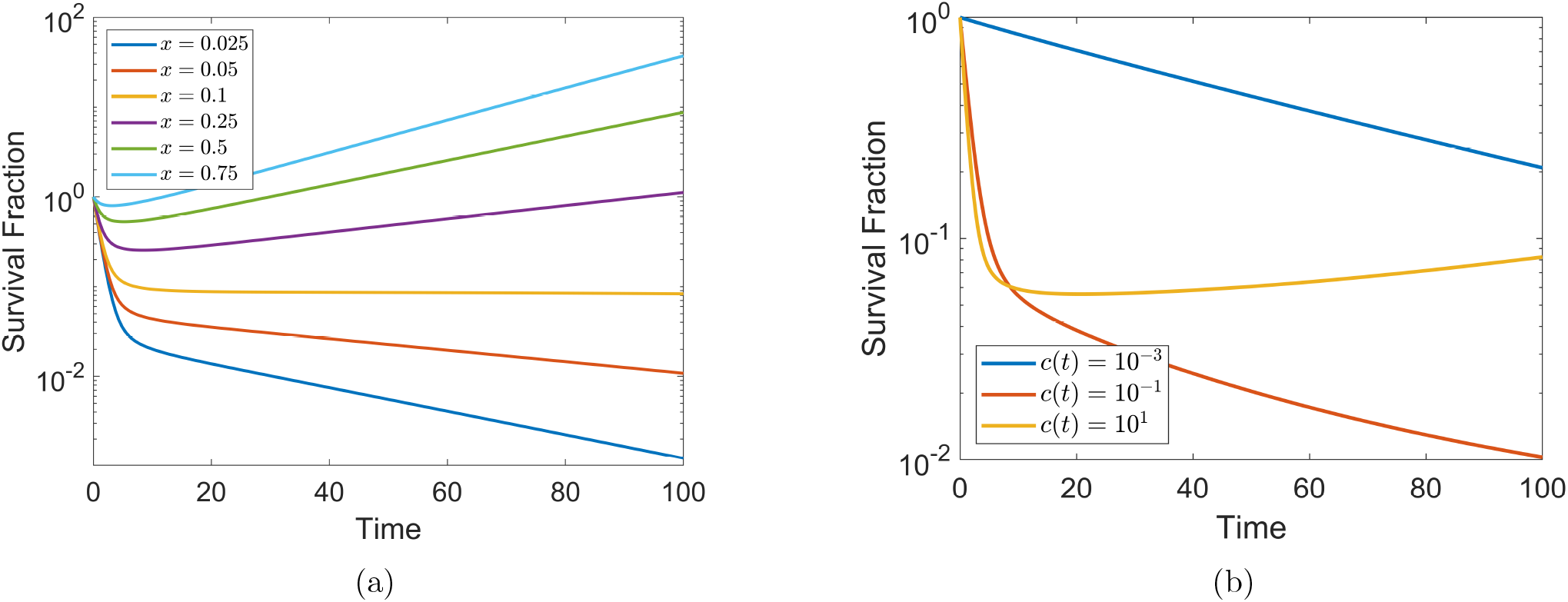
(a) Survival curves for single-cell clones with different *x* values (ctp) throughout the treatment with *c*(*t*) = 1. (b) The total population curves at various drug concentrations (*c*(*t*)).

The first set of simulations with *c*(*t*) = 1 reveals that subpopulations with different CTPs asymptotically converge to one of two outcomes: gradual extinction or cycling persistence (Figure 2(a)). When the size of each *x*-clone (i.e. a single-cell clone with a given *x*) is weighted by the initial CTP distribution (that is, the initial probability density in *x* before treatment), the total population ultimately begins to regrow, albeit at a relatively slow rate (Figure 2(b)). This behavior reflects the near extinction of low-CTP cells, which initially dominated the population distribution, followed by the emergence of cycling persistence among survivors through advection in the *s*-phenotypic space. Increasing the drug concentration enhances the initial killing rate, but also promotes extensive phenotypic adaptations, potentially leading to cycling persistence (Figure 2(b)).

Conversely, reducing *c*(*t*) excessively can result in insufficient killing and suboptimal long-term outcomes (Figure 2(b)). It is important to note that during most *in vitro* drug-dose response experiments, viability is measured 1-9 days after the initiation of the treatment. As shown in Figure 2(a), cells with low *x* values are predestined for extinction, whereas cells with sufficiently high *x* values may regain the ability to cycle. This suggests the existence of a critical CTP value, *x*_*c*_, for a given constant drug concentration, such that a time at which 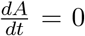 exists for *x > x*_*c*_ but not for *x < x*_*c*_ (Figure 3(a)). In this context, determining is equivalent to finding *x* that makes the asymptotic value of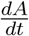 zero (Figure 3(b)). With parameters chosen for the simulation (see Table S1), we numerically estimated *x*_*c*_ ≈ 0.103 for *c*(*t*) = 1.

**Figure 3:**
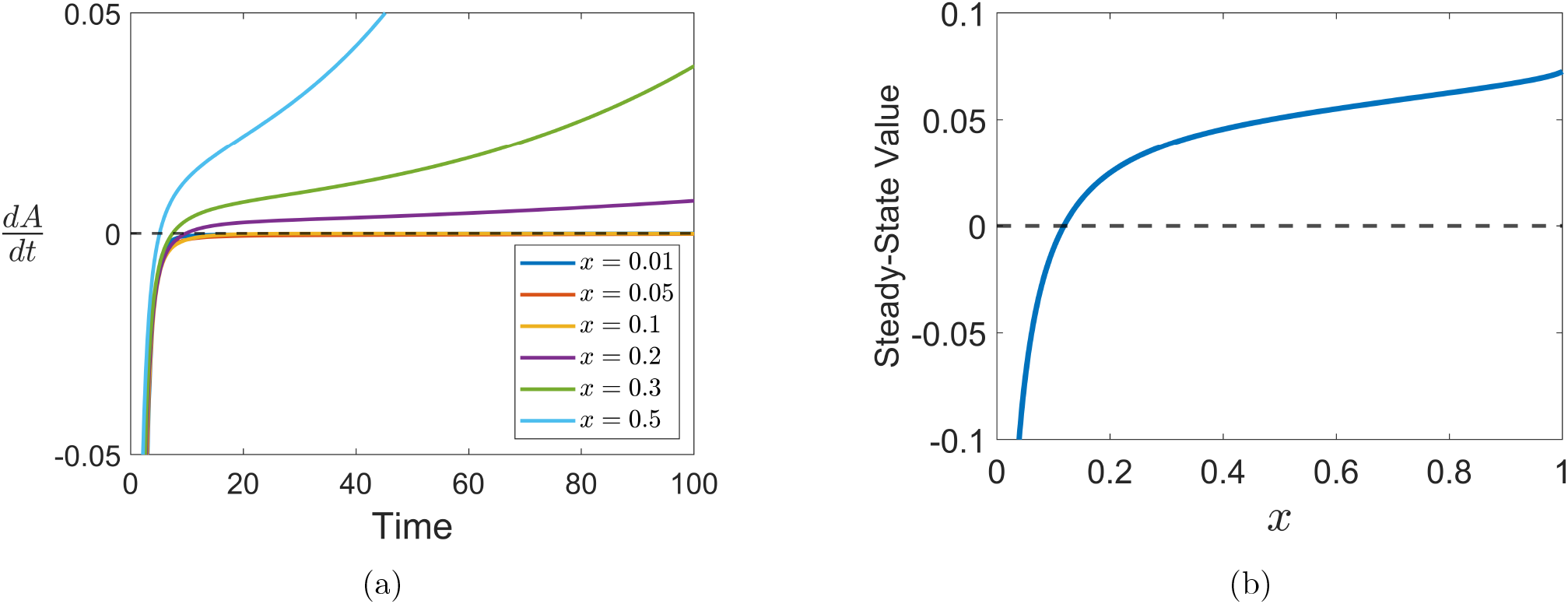
Plots of 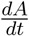for different values of *x* (CTP) and (b) the asymptotic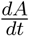 as a function of *x* for *c*(*t*) = 1.

We show that, under certain assumptions, *x*_*c*_ can be approximated through a more analytical approach. Briefly, our method assumes that the advection strength can be reasonably approximated as some constant value (*v*_*c*_), and that the net growth rate (*r* − *k*) vanishes at the steady-state limit. Then, the resulting simpler Ornstein-Uhlenbeck equation is solved using the Green function (see Materials and Methods for more details). For example, with *c*(*t*) = 1, our approach yielded *x*_*c*_ = 0.102, which is very close to the critical value determined by manually solving the fully time-dependent PDE (*x*_*c*_ = 0.103). While this approximation method does not yield exact values of *x*_*c*_ for every *c*_0_, it still qualitatively replicates the correct behavior (Figure 4).

**Figure 4:**
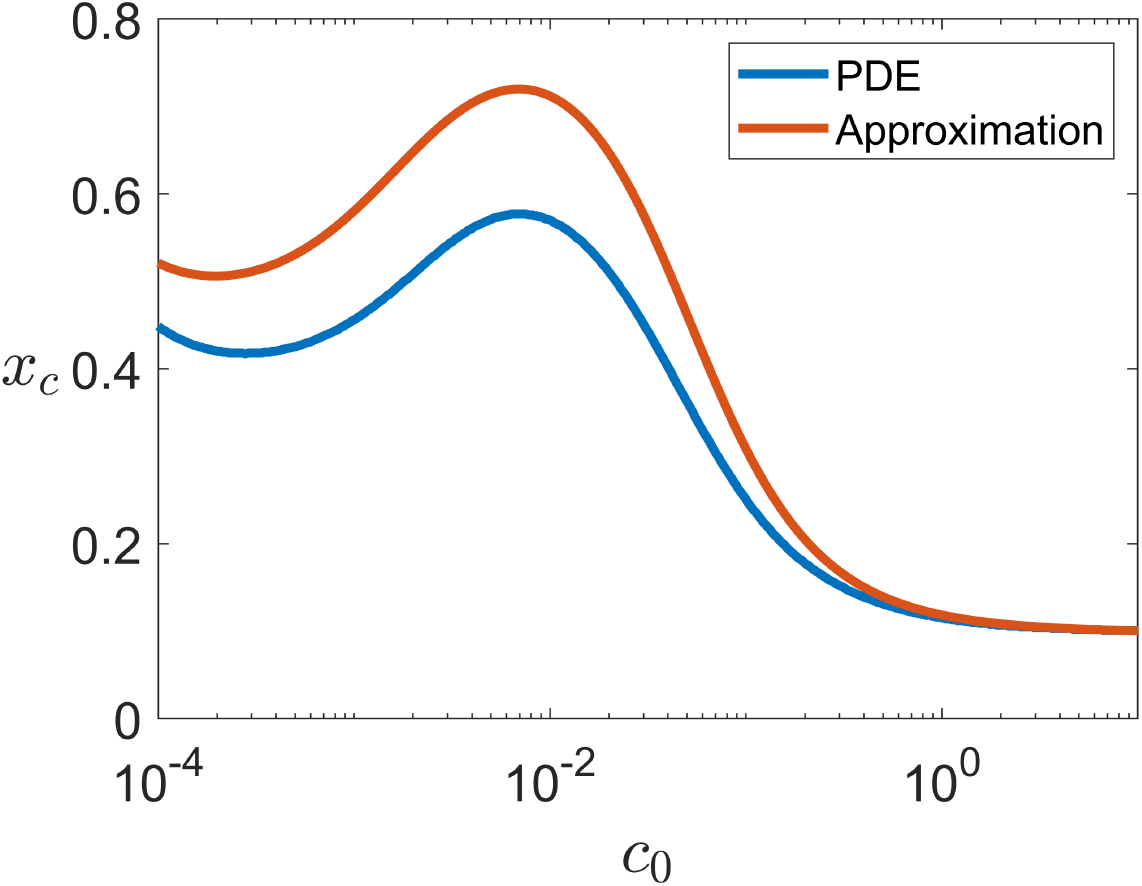
Values of critical CTP (*x*_*c*_) calculated for different *c*(*t*) = *c*_0_ using our approximation method and by solving the PDE.

Determining *x*_*c*_ allows us to predict the fate of single-cell clones with a given CTP, analogous to the experimental work by Oren et al. [17], where different lineages of PC9 cells were treated with osimertinib, an anti-EGFR TKI, for 14 days and subsequently classified as “drug sensitive”, “persisters with small progeny”, and “persisters colonies”. In our case, “persister with small progeny” refer to DTPs with minimal or no cell division while “persisters colonies” denote cycling persisters. Therefore, the fraction of the population likely to attain cycling persistence can be estimated using an initial CTP distribution. Finding *x*_*c*_ can also aid in optimizing therapeutic doses. Selecting a dose with a high *x*_*c*_ reduces the likelihood of cycling persistence while enhancing cell killing. Our method suggests that the optimal constant dose yielding the maximum *x*_*c*_ is *c*_0_ = 6.9 × 10^*−*3^, while solving the PDE yields *c*_0_ = 6.3 × 10^*−*3^ (Figure 4). This condition is consistent with the data presented in Figure 2(b), where the most effective concentration is seen to be between 10^*−*3^ and 10^*−*2^.

### 2.3 Drug Holidays

A potential weakness of continuous therapy, as demonstrated by previous studies [33, 34] and real-world clinical data [35, 36], as well as our simulations from the previous section, is the induction of DTPs and cycling persisters, which may acquire resistance after a prolonged survival, ultimately leading to treatment failure. Clinicians have long observed the emergence of acquired persistence and resistance due to inadvertent adaptation mechanisms and the unintended selection pressures caused by cytotoxic therapeutics [37, 38].

In response, concepts such as drug holidays [32, 39, 40] and adaptive therapy [41] are gaining traction and being increasingly explored in clinical settings. The rationale behind drug holidays is that they allow more sensitive cells to repopulate and occupy a larger fraction of the tumor population during periods of drug withdrawal, instead of continuously administering maximally tolerated dose (MTD) or other high doses to maximize short-term cell death. Promising results from early clinical trials, particularly in prostate cancer, suggest that adaptive therapies with intermittent drug holidays can delay disease progression more effectively than continuous treatment [42]. Mathematical models have also been developed to predict the potential benefit of adaptive therapy [43]. However, most of these models consider two discrete states – sensitive and resistant – without accounting for the continuous phenotypic transitions observed in persistence [44].

Using our continuous model, we simulated various treatment regimens incorporating drug holidays over a total duration of 100 days to investigate their impact on phenotypic adaptations and overall cell survival. These results were compared with the continuous constant-dose administrations at *c*(*t*) = *c*_0_. For all simulations, the total drug exposure was constrained to match the area under the curve (AUC) of the respective continuous treatment, *c*_0_*T*, where *T* = 100 days represents the total therapy duration.

During the “drug-on” period (*c*(*t*) *>* 0), cells adapt to drug-induced stress, primarily through *s*-dynamics. In contrast, during the drug holiday phases (*c*(*t*) = 0), the population seeks to relax toward its initial steady-state distributions in both *s* and *x*. These relaxation dynamics reduce fitness in the phenotypic space by reversing drug-induced advection in *s* and increasing the prevalence of cells with lower CTP (*x*). For simplicity, the drug-on and holiday durations were kept equal, implying higher drug-on doses compared to the continuous regimen to achieve the equivalent drug exposure.

Simulations revealed that the efficacy of drug holidays heavily depends on the rates of global growth (*r*_*x*_) toward the carrying capacity (*N*_*ss*_) and epigenetic diffusion (*µ*). A rapid growth in the absence of the drug can potentially neutralize the expected benefit from the holiday therapy. On the other hand, higher values of *µ*, enable the population to recover low-CTP cells more effectively during holiday periods, leading to greater tumor reduction when the drug is applied. For instance, with two holidays and *c*_0_ = 1, higher *r*_*x*_ values consistently resulted in worse outcomes than the continuous regimen (Figure 5(a)). Conversely, simulations with lower *r*_*x*_ values consistently predicted more favorable outcomes than the continuous case (Figure 5(a)). Simultaneously, higher *µ* values resulted in better outcomes, while lower *µ* values slowed the relaxation toward the initial distribution in *x*, diminishing the benefits of drug holidays (Figure 5(b)). The latter result suggests that drug holidays may be particularly effective against tumors with a more “malleable” epigenetic landscape.

**Figure 5:**
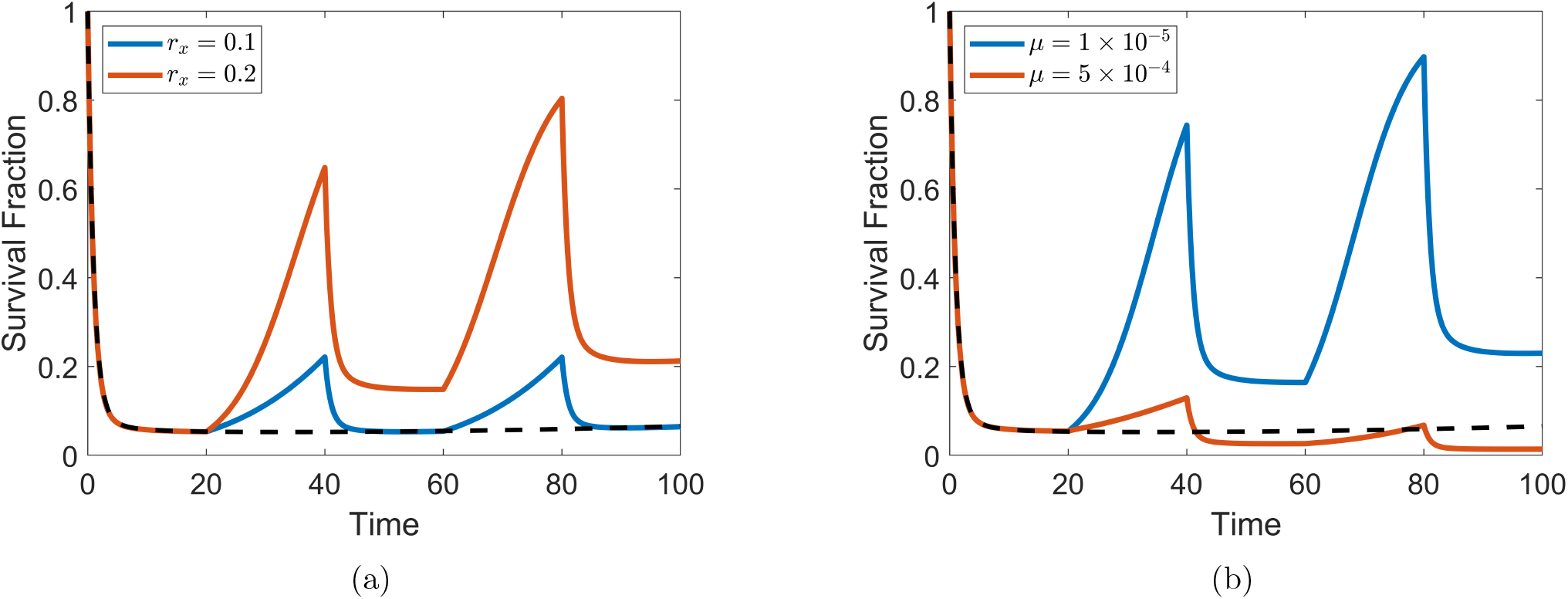
The peaks and troughs of the total tumor population during treatment with two drug holidays compared to the curve during continuous therapy with *c*_0_ = 1 with different (a) *r*_*x*_ values (*µ* = 3.6 × 10^*−*5^) and (b) *µ* values (*r*_*x*_ = 0.2).

Additional simulations examined the effect of varying the number of holidays (*n*) while keeping total exposure fixed. In these simulations, *µ* and *c*_0_ were fixed at 3.6×10^*−*5^ and 1, respectively. The results interestingly revealed that arbitrarily increasing the number of holiday leads to worse outcome (enhanced tumor cell survival) compared to the AUC-equivalent continuous treatment for tumors with high *r*_*x*_ (Figure 6(a)). Although the case with lower *r*_*x*_ yielded more promising outcomes, arbitrarily increasing the number of holidays eventually produces diminishing returns (Figure 6(a)). With more holidays, the drug-on doses are higher (to preserve AUC equivalence), intensifying selection pressure and phenotypic adaptations. At the same time, shorter holiday intervals provide insufficient time for the population to revert to initial distributions. In fact, even a single holiday is sufficient to shift the CTP distribution further to the right compared to the continuous treatment (Figure 6(b)). Of course, implementing excessively many holidays within a 90-day treatment period is neither feasible nor practical in clinical settings, but the simulations provide mathematical confirmation of our intuitive expectations. Overall, these findings underscore the importance of carefully balancing drug-on periods, which exert cytotoxicity but also drive adaptation and selection, with holiday durations sufficient to allow the repopulation of less fit and low-CTP cells.

**Figure 6:**
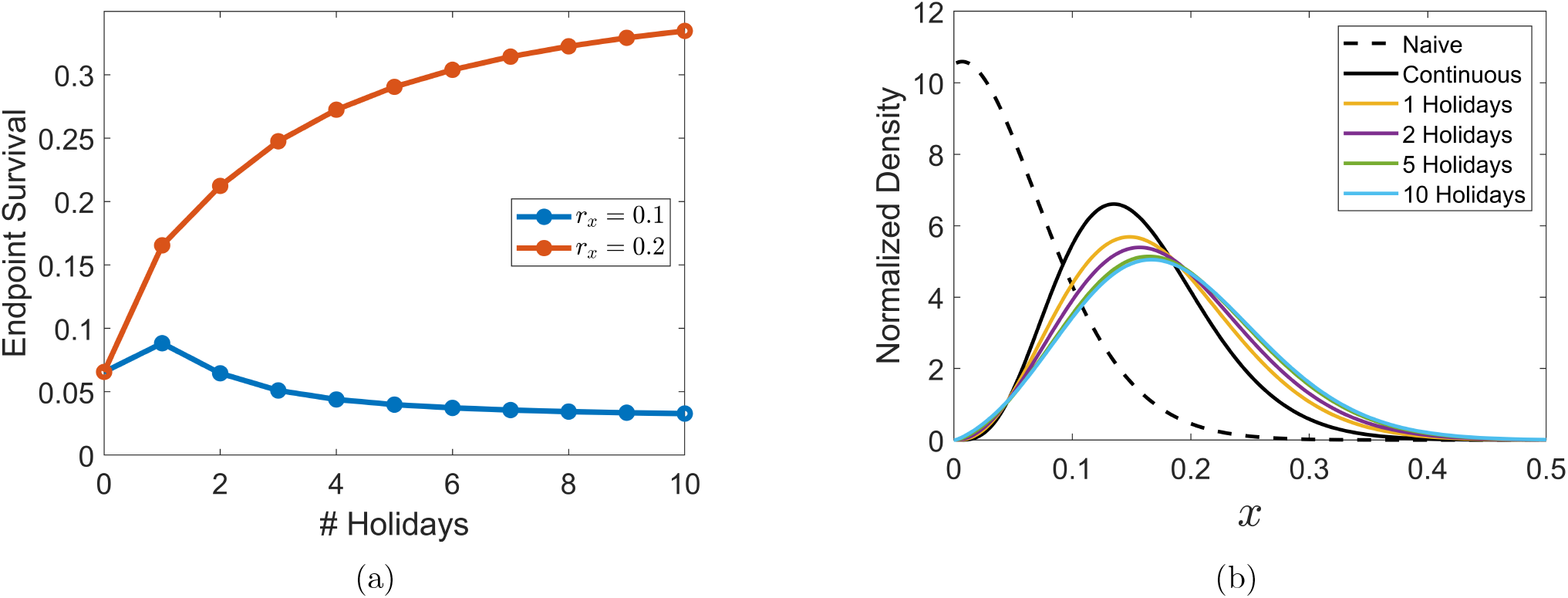
(a) The overall surviving fractions of tumor population and (b) the CTP (*x*) distribution after *T* = 100 days of treatment with a fixed total dosage but different numbers of holidays.

In addition to holiday frequency, the outcomes also depended on the drug dose (*c*_0_). With sufficiently low *r*_*x*_, simulations with two holidays (*n* = 2) and varying *c*_0_ revealed a nuanced relationship between dose and the efficacy of holidays, as illustrated in Figure 7. At low doses, drug holidays were ineffective and offered no real benefit compared to continuous treatment (Figure 7). However, at sufficiently high doses, capable of inducing significant drug tolerance over longer time scales, the two drug holidays improved the treatment outcome at day 100 (Figure 7). Together with other simulations mentioned earlier, this finding highlights the complex interplay between tumor plasticity, treatment schedules, and drug dosing. In particular, optimizing drug holiday regimens requires consideration of not only total drug exposure and timing but also the pharmacokinetics/pharmacodynamics (PK/PD) of the specific drug and the adaptive capacity of the tumor population.

**Figure 7:**
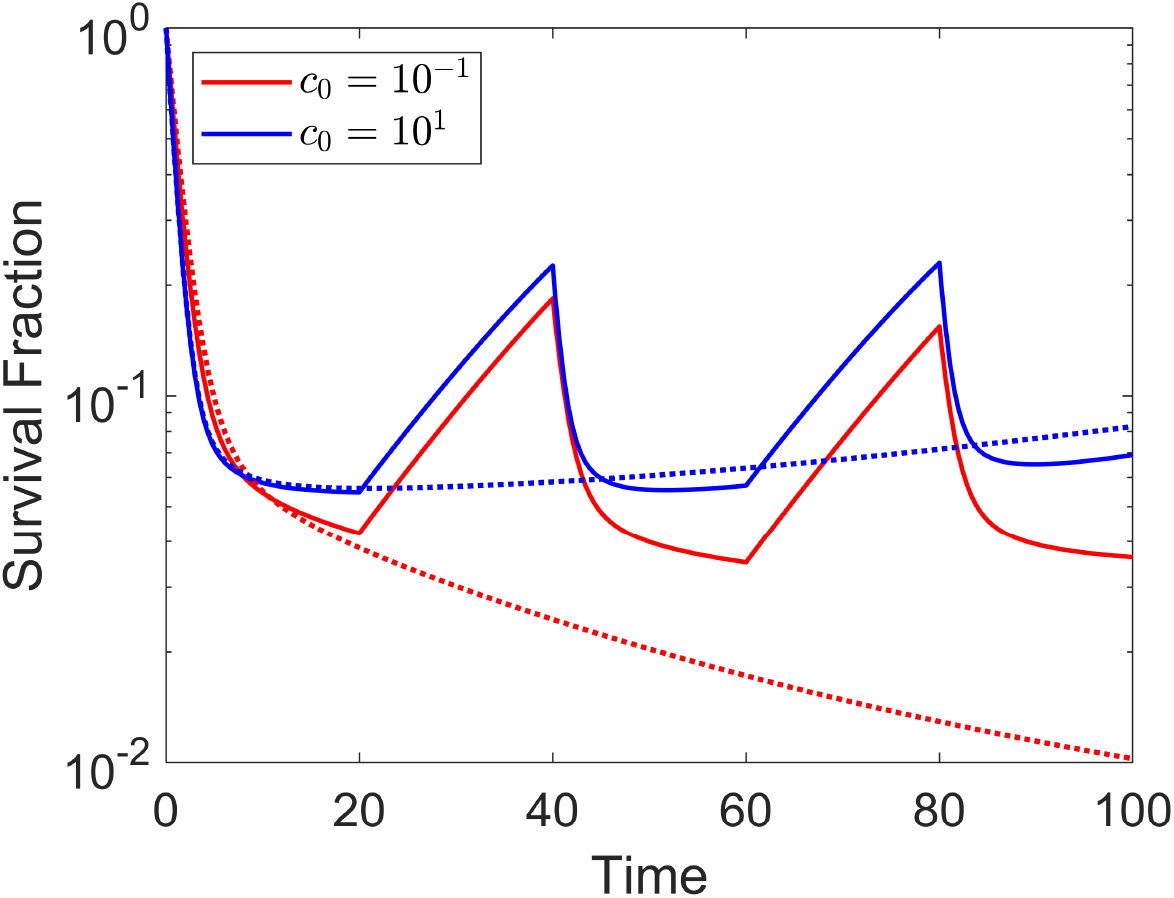
Simulated total population curves for both continuous (dotted lines) and two-holiday therapies (solid lines) with different reference doses (*c*_0_).

### 2.4 The Continuum of Drug Tolerance

As mentioned before, the populations of sensitive and drug-tolerant/resistant cells are often viewed as discrete states, dynamics of which are governed by a system of ODEs [46, 47]. In contrast, our theoretical model assumes phenotypic transitions across a continuous spectrum of cell states. Both approaches seem to generate a sufficiently precise fit to experimental data and accurate predictions, at least in near steady-state limits. In fact, one could even argue that adding intermediate states between “sensitive” and “resistant” states in the model does not significantly improve fits to data [48].

Interestingly, recent dose escalation experiments performed by França et al. [45] suggest the existence of “resistance continuum” along which tumor cells can adapt to and tolerate therapies. Briefly, the group initially exposed a population of cancer cells to a continuous low-dose treatment with a cytotoxic agent.

After confluence was reached, a small fraction was passaged and then, this time, was exposed to a higher dose (e.g. doubled) of the same drug. This was repeated for multiple (6-9 times, depending on the cell line) iterations, generating a slightly more resistant batch than the previous cycle at each step. The authors performed this experiment on 5 different cell lines derived from ovarian, lung, and melanoma cancers.

Although the authors were less interested in the characterization of the gradual adaptation dynamics and, instead, were primarily concerned with generating populations of cells with varying degrees of resistance to identify metabolic and chromatin-related differences between them, we extrapolated the “time to confluence” (the number of days to reach confluence at each cycle) from their work. We then compared these values with those obtained from the simulations of their dose-escalation experiment using our mathematical model. Without attempting to meticulously fit the model to their data, our simulations did not yield exactly the same values but successfully predicted the observed overall behaviors of the cell lines (Figure 8 and also see Figure S1 and Table S2 in Supporting Information). This result not only corroborates our PDE approach, but also highlights the sheer diversity in the degree of drug-tolerance/resistance in tumors resulting from extensive cellular phenotypic plasticity under stress.

**Figure 8:**
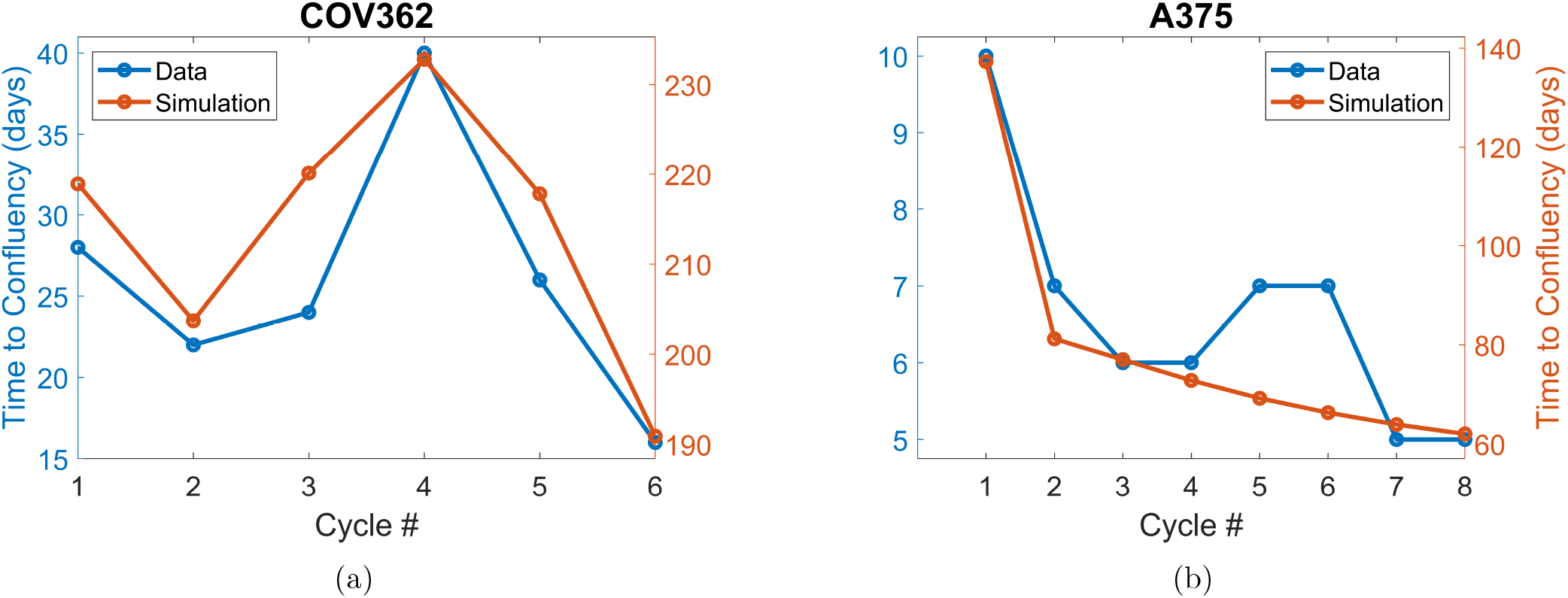
Times to confluency extracted from França et al. [45] (blue) and calculated with our simulation (orange) for (a) COV362 high-grade serous ovarian cancer and (b) A375 melanoma cell lines.

## 3 Discussion

Drug persistence, as distinct from resistance, is receiving growing attention among clinicians and cancer biologists. Previous efforts to experimentally probe and theoretically model the phenotypic plasticity of tumor cells, which leads to the emergence of DTPs and cycling persisters, confirm the reversibility of the phenomenon, albeit with a likely significant time lag [14, 20, 27]. These studies suggest that persistence is likely to be drug-induced at least in cancer systems. Motivated by these findings, we developed a mathematical model to describe cancer cells as a population distributed across a probability density in an abstract space representing epigenetic (*x*) and phenotypic (*s*) degrees of freedom. This model characterizes how drug exposure enables certain cells to withstand cytotoxicity, enter a quiescent state, and even restore the ability to cycle and proliferate. In our theoretical framework, the presence of drug drives the cells to advect in *s*, gaining fitness and tolerance to the drug’s pharmacological effect. Further advection to higher *s* may lead to cycling persistence. Conversely, in the absence of the drug, the population gradually regains drug sensitivity by reversing the advection in *s* and approaching the initial steady-state distribution in *x*.

Overall, our model captures several key characteristics and properties of persistence. First, it does not presume the existence of persistent cells prior to treatment. However, different cells have varying probabilities of becoming persistent, represented by variable *x* (CTP), which we consider epigenetically determined [49, 50]. The fate of these cells depends on their initial positions in *s*, advection rate (*v*(*s, c*(*t*))), and the net proliferation/death rate (*r*(*x, s, c*(*t*))−*k*(*s, c*(*t*))). Second, our model accounts for the reversibility and allows cells to reclaim the sensitivity once the drug is removed. This reversal occurs through two mechanisms: (a) the cessation of advection, effectively removing *v* from the PDE, and (b) the gradual return of the *x*-distribution to its steady-state form. However, this process can be slow, especially for the *x*-dynamics, if *µ* is small, making it difficult to observe substantial changes unless drug holidays are sufficiently long. Lastly, the model takes into account the possible emergence of cycling persistence. Importantly, the initial quiescence of persistent cells does not result solely from the balance between birth and death rates. Experimental evidence confirms the existence of states resembling actual cell-cycle arrest among DTPs [32]. Our model enables a subset of DTPs to transition into cycling persistence by advecting to higher *s* values, where *R*_*s*_(*s, t*), initially nearly zero, becomes sufficiently large to support net positive growth.

Simulations of various treatment schedules elucidates interesting and critical factors to consider when designing and optimizing therapeutic interventions. Under continuous treatment with a constant concentration (*c*(*t*) = *c*_0_), our model predicts the existence of a critical CTP, *x*_*c*_, that can be used to determine the fates of single-cell clones – cycling persistence or gradual extinction. We demonstrated that our semianalytical method for approximating *x*_*c*_ produces results qualitatively align with those obtained by fully time-dependent PDE. Moreover, calculating *x*_*c*_ for different *c*_0_ enables us to identify optimal continuous dose that maximizes the CTP threshold for cycling persistence. We also showed drug holidays could mitigate the induction of (cycling) persistence by allowing cells to revert toward *s*_0_(*x*) in *s* and repopulating cells with low CTP. However, several crucial caveats must be noted. First, drug holidays were only effective for cells with small *r*_*x*_, the growth rate toward the carrying capacity in the absence of the drug. Second, the efficacy of holidays was significantly enhanced in populations with high *µ*, suggesting that epigenetic “malleability” plays a crucial role in determining treatment outcomes under intermittent regimens. Furthermore, arbitrarily increasing the number of holidays resulted in diminishing returns (for cases with sufficiently small *r*_*x*_), accompanied by a shift in the distribution at the treatment endpoint toward higher *x* values. Lastly, the benefits of drug holidays were only apparent (once again, for sufficiently small *r*_*x*_) at sufficiently high baseline doses, emphasizing the need for dose optimization.

Increasing the drug dose enhances the killing rate but may simultaneously induce a greater number of DTPs and cycling persisters over longer timescale. Finding an optimal balance between eliminating sensitive cells and minimizing the generation of persisters is crucial for designing treatment algorithms that maximize therapeutic efficacy and reduce the risk of a limited response. Additionally, adverse effects and potential toxicities must also be considered especially *in vivo* and clinical settings. Combining our theoretical work with available PK/PD data has the potential to facilitate highly efficacious and safe therpay optimization.

Surprisingly, there is limited experimental and theoretical literature that frames persistence as a continuous spectrum of phenotypic traits or explores how varying drug concentrations and administration schedules influence the induction of these traits. The conventional binary categorization of cell states into either “sensitive” and “resistant” (or “persistent”) along with ODEs to model transition kinetics between these discrete states offers computational simplicity and tractability. Also, the introduction of intermediate states between the binary extremes may not significantly improve model performance in terms of data fitting [48]. However, this approach may obscure the underlying phenotypic heterogeneity and prompts an uncomfortable but important philosophical question: which modeling approach offers a higher degree of isomorphism to biological reality and/or is more useful for practical usage? It is, therefore, encouraging to see studies like França et al. [45], which exposed cells to a range of drug concentrations to generate populations with varying degrees of tolerance to cytotoxicity, thereby effectively demonstrating the existence of a phenotypic continuum through which tumor cells adapt to therapy.

Similarly, Berger et al. [27], which successfully produced numerous single-cell clones characterized by distinct continuous trait parameters (CTP or *x*), further supports the view that durg-tolerance phenotypes span a spectrum rather than fall into discrete categories. Together these experimental findings lend strong support to our modeling framework, which treats adaptation in persister populations as a continuous process. Nonetheless, further experimental validation – particularly studies comparing the response of distinct lineages within genotypically identical populations across varying drug concentrations – is essential to fully substantiate our theoretical approach.

Our model, however, operates under several assumptions and has certain limitations. For example, our phenotypic space governs both cell survivability and proliferation, as *r* and *k* are both functions of *s*. As a result, DTPs are treated as a required precursor state before cells can transition into cycling persisters. This approach differs from models proposed by Clairambault et al. [31], which separates suvivability and the ability to proliferate into two distinct phenotypic dimensions, but is consistent with the experimental observation by Sharma et al. [14]. Moreover, the physiological mechanisms underlying our simulation results derived from an abstract mathematical remain uncertain. We did aim to treat persistence as a pharmacologically driven and drug concentration-dependent process, using functional forms grounded and commonly applied in biological studies. One promising future direction is to investigate the complex metabolic network and its dynamics in cancer. Significant differences in metabolic states among cells with varying levels of drug tolerance and resistance have been identified by multiple groups [9, 51, 52].

In summary, we present a mathematical model that captures the induction of DTPs and cycling persisters in a drug concentration-dependent manner, aligning with various trends and properties in experimental and clinical data. As discussed, mapping relevant biochemical and metabolic processes to the outcome predicted by the model would further confirm our theory. Efforts like ours could significantly advance the understanding of persistence and acquired resistance, helping clinicians design optimal personalized treatments for cancer patients. Moreover, we believe that our mathematical approach can be generalized to characterize not only how tumor cells survive treatment but also how other organisms adapt to stimuli and stress beyond cytotoxic agents. It is well-documented that bacteria and yeasts, for example, exhibit remarkable cellular plasticity under diverse forms of stress, with these adaptations often enhancing their ability to respond to entirely different challenges, including even the development of persistence against anti-microbial treatments [8, 21, 53, 54]. Our framework may offer a promising tool for investigating the interplay adaptation mechanisms for various stress factors and how they are coupled to collectively endow cells with survival advantages under different conditions. Ultimately, we aim to continue developing robust theoretical models that predict tumor responses to treatment and propose strategies to combat cancer cells by disrupting their adaptability under drug exposure.

## 4 Materials and Methods

All numerical simulations, as well as solving partial differential equations, and figure generations were performed using MATLAB R2022b (MathWorks).

### 4.1 The Model

The population distribution across *x* ∈ (0, 1) and *s* ∈ (−∞, ∞) obeys the dynamics described by the following PDE:

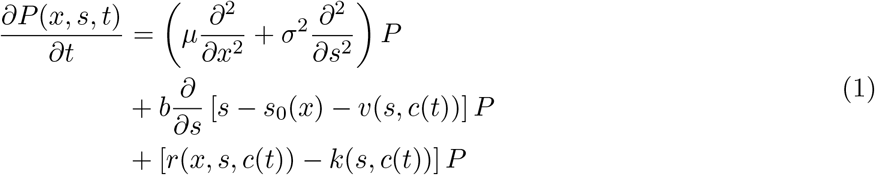

where *c*(*t*) is the drug concentration as a function of time. The population is subjected to diffusion in both *x* and *s* with *µ* and *σ*^2^ as the respective transition rates.

The advection term has the generic forms

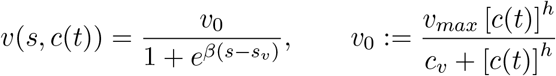

where *β, s*_*v*_, and *c*_*v*_ are parameters that modulate the functional shape of *v*(*s, c*(*t*)). Similarly, the death rate is chosen to be

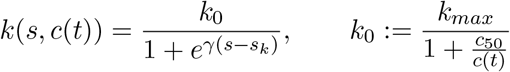

where *γ* and *s*_*k*_ are both constants that determine the shape and steepness of *k*(*s, c*(*t*)).

Lastly, the proliferation term, *r*(*x, s, c*(*t*)), is assumed to be:

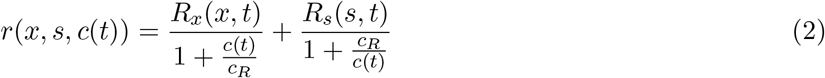

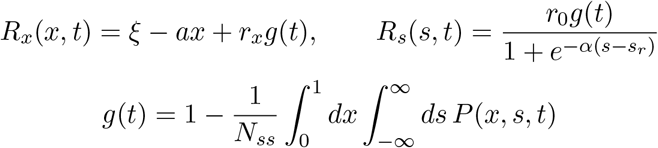

Note that *g*(*t*) represents the logistical damping of growth as the population reaches its carrying capacity *N*_*ss*_. Notably, *r*(*x, s, c*(*t*)) becomes nearly completely dependent on *x* when *c*(*t*)*/c*_*R*_ → 0, while it is predominantly influenced by *s* when *c*(*t*)*/c*_*R*_ → ∞. With a sufficiently small *c*_*R*_, Equation 2 can be reasonably approximated as:

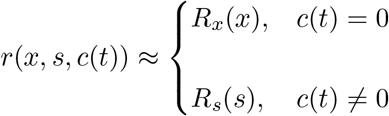

We assume that a sufficiently long time has elapsed prior to treatment such that the initial population distribution follows the steady-state solution of Equation 1 in the absence of drug (*c*(*t*) = 0). *P* can be decomposed into a steady distribution along *x* and a conditional probability density along *s* for each given *x*. That is, at *t* = 0, we can express 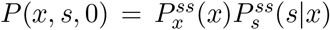. Under this assumption, 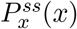 and 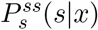 satisfy the following ODEs with *g*(*t*) = 0:

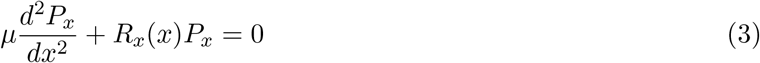

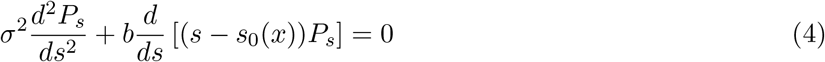

The solutions to Equation 3 and 4 are an Airy distribution with 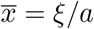(Equation 5) and a Gaussian distribution centered around *s*_0_(*x*) (Equation 6), respectively, where 𝒩 is a scaling constant such that 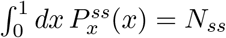.

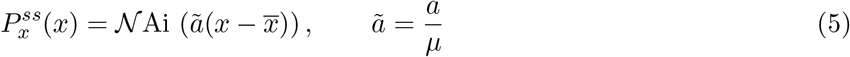

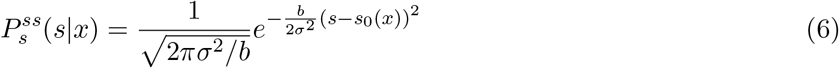

It is worth noting that the motion of these tumor cell “particles” at the two extremes of the piegenetic space (*x* = 0 and *x* = 1) is unclear. Our operational definition of CTP does not allow *x* to be defined outside of the interval (0, 1), and the exact characterization may vary depending on the chosen boundary condition. However, we expect that Equation 5 captures behavior away from the boundaries reasonably well. To confirm our intuition, we also performed a stochastic simulation, which revealed that the normalized distribution of 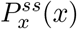 closely follows the sharp Airy-like profile with a peak at some low *x* value (see Figure S2 in Supporting Information), consistent with Equation 5. Meanwhile, *s*_0_(*x*) is self-consistently determined from *x* in accordance with the operational definition of CTP:

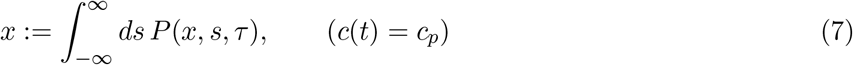

Based on Equation 6 and 7, having a larger *x* value bestows a single-cell clone with the initial distribution in *s* with a higher mean phenotype.

We assume that phenotypic transitions occur at a much faster timescale than epigenetic changes. This allows us to impose the condition *µ* ≪ *σ*^2^. This implies that the presence of the drug (*c*(*t*) *>* 0) forces the dynamics to be primarily *s*-dependent, while in the absence of the drug (*c*(*t*) = 0), the *x*-dynamics play a more noticeable role. Under these assumptions and for cases where *c*(*t*) is either some positive constant or zero (e.g. similar to an *in vitro* experiment), Equation 1 can be approximated as:

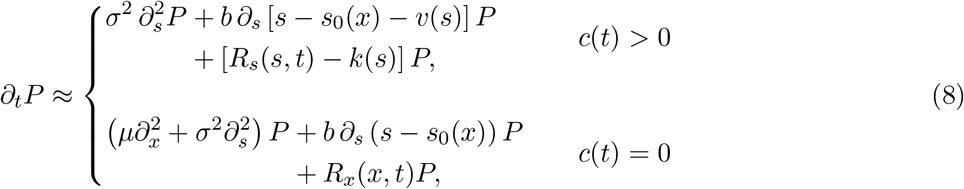

When *c*(*t*) = 0, the solution of Equation 8 will relax to the initial conditions i.e. Airy distribution in *x* and Gaussian distribution in *s*, as described in Equation 5 and 6, respectively. When the drug is present (*c*(*t*) *>* 0), the advection rate *v*(*s*), birth rate *R*_*s*_(*s, t*), and death rate *k*(*s*) depend solely on the phenotypic state *s*. Consequently, in the presence of the drug, the population initially declines, but the probability density shifts toward higher *s* values, enabling the cells to acquire enhanced tolerance to the treatment. If a subpopulation starts with a very high *s*_0_(*x*) and/or is advected sufficiently in the positive *s*-direction, these cells have the potential to become cycling persisters.

Treatments can be accurately simulated, and the outcomes can be estimated using Equation 8.

### 4.2 Continuous Dose Simulation

Unless noted otherwise or to have been varied, all simulations were performed using the parameters with values shown in Table S1.

### 4.3 Survival Curves

The survival curves for a given *x*-clone can be computed and plotted using:

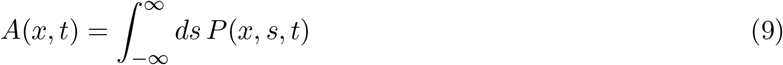

Similarly, the total population curve of tumor cells at any time point can be determined using:

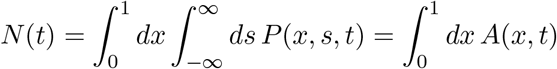

### 4.4 Calculation of Critical CTP

During a constant-dose continuous treatment, we only have to solve the *c*(*t*) *>* 0 case of Equation 8.

In the near steady-state limit, we assume that we can make the following assumptions:

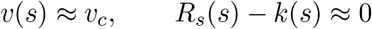

where *v*_*c*_ is some constant, of course, this approximation breaks down on a short timescale or when |*r* − *k*| becomes significant. However, at near steady-state regimes, where the population experiences minimal change, we believe these assumptions are qualitatively valid. Under these conditions, Equation 8 further simplifies to:

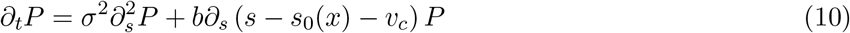

Equation 10 can be solved by integrating the product of its initial steady-state solution, 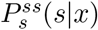, and the Green’s function *G*(*s, s*^*′*^, *t, t*^*′*^|*x*) for the Ornstein-Uhlenbeck process given by:

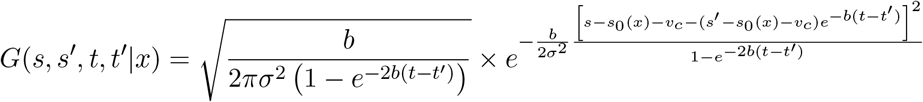

Thus, the full solution for *P* (*x, s, t*) in this case is:

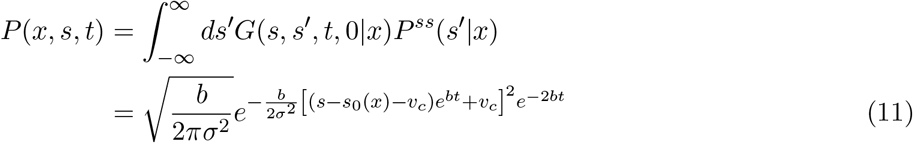

Meanwhile, from Equation 9 and the *c*(*t*) *>* 0 case of Equation 8, one can easily derive:

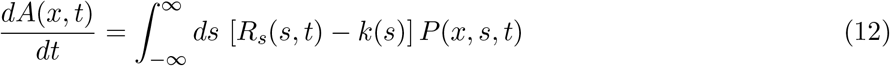

By substituting Equation 11 into Equation 12, we can obtain an explicit expression for *dA/dt* at steadystate. This allows us to plot the curve and determine the critical value *x*_*c*_, where a root exists for *x > x*_*c*_, but does not for *x < x*_*c*_. However, it suffices to focus on the asymptotic behavior at steady-state. At *t* → ∞, the existence of a root of Equation 12 is determined entirely by *s*_0_(*x*):

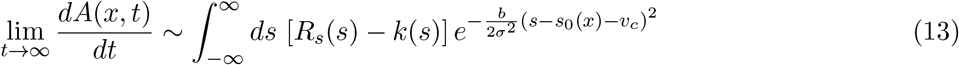

For a given constant *c*(*t*) = *c*_0_, we chose *v*_*c*_ = *v*(*s*_*rk*_), such that at *s* = *s*_*rk*_ and *g*(*t*) = 1, *R*_*s*_ −*k* = 0. Since *dA/dt <* 0 at small *t* for all *x <* 1, the critical value *x*_*c*_ precisely corresponds to the root of Equation 13.

### 4.5 Drug Holiday Simulations

Once again, unless varied, all nominal values of the parameters used in the simulations can be found in Table S1.

To simulate treatment regimens with drug holidays, we now have to take account of both cases of Equation 8. Under a regimen with *n* isochronal drug holidays between *t* = 0 and *t* = *T* (beginning and ending the treatment with drug-on periods), *c*(*t*) is given by:

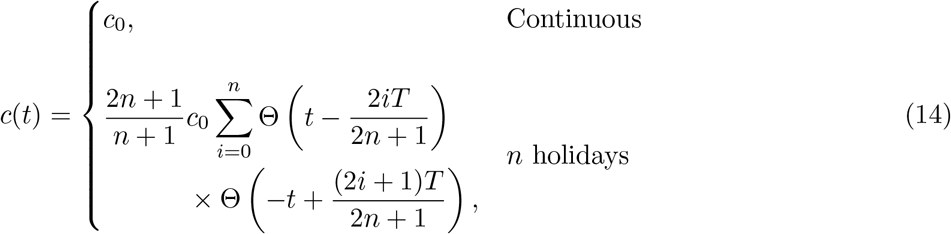

Here, Θ(*y*) is the Heaviside step function, which yields 0 for *y <* 0 and 1 for *y >* 0. Note that Equation 14 ensures that the AUC, 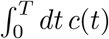is fixed for both cases.

### 4.6 Dose Escalation Simulation

To investigate the continuous spectral nature of drug-tolerant states, we performed simulations of the dose escalation experiment conducted by França et al. [45]. The times to confluence for each cell line were extrapolated by simply subtracting the two consecutive cumulative days reported from their experiments. For the simulations, equivalent initial doses were first approximately determined for each cell line based on the viability data provided by the authors. For example, the initial dose for Kuramochi was estimated to be that corresponding to ∼ 0.95 survivability after 9 days of treatment.

We then replicated the dose escalation scheme by increasing the dose at each cycle by exactly the same proportion. Each cycle during the simulation lasted until the confluence was reached. In accordance with the common laboratory practice, we defined confluence to be approximately 0.7 × *N*_*ss*_. However, during the correspondence, the authors indicated that the cells undergo morphological changes while becoming increasingly drug resistant. For example, if 10^6^ cells were initially passaged and the threshold for confluence was initially 5 million cells, by the end of 9 cycles with Kuramochi cells, for example, the population at confluence increases to 7-10 million cells. In our simulations, we took this into account by linearly increasing the confluence threshold from 5 million at the start of the experiment to 10 million by the 9th cycle (however, the 5 cell lines were subjected to different numbers of cycles). Using the days to confluence data from our simulation, we manually adjusted a few of the parameters (Table S2) to replicate qualitative trends observed by França et al. [45].

## Acknowledgements

The authors acknowledge the support of the NSF through the Center for Theoretical Biological Physics, grant no. PHY-2019745. We would also like to thank Gustavo S. França for corresponding with us and providing us with insights on the dose escalation experiment [45].

