## Supporting Information for "Modeling a Continuum of Drug-induced Persistence during Targeted Cancer Therapy"

### 1 Parameter Values for Continuous and Holiday Treatment Simulations

All continuous and holiday treatment simulations were performed using the following nominal values of parameters (Table 1) unless otherwise noted in the main text.

Table 1: Parameters used in Treatment Simulations

| Parameter | Value | Parameter | Value |
|---|---|---|---|
| $\mu$ | $3.6 \times 10^{-5}$ | $\gamma$ | 4 |
| $\sigma$ | 0.3 | $s_k$ | 0 |
| $b$ | 0.09 | $r_0$ | 0.08 |
| $v_{\max}$ | 10 | $\alpha$ | 1 |
| $h$ | 1 | $s_r$ | 2.5 |
| $c_v$ | 0.05 | $\xi$ | 0.012 |
| $\beta$ | 0.5 | $r_x$ | 0.2 |
| $s_v$ | 0 | $a$ | 0.18 |
| $k_{\max}$ | 1 | $c_R$ | $1 \times 10^{-6}$ |
| $c_{50}$ | 0.05 | | |

### 2 Simulation of Dose Escalation Experiments

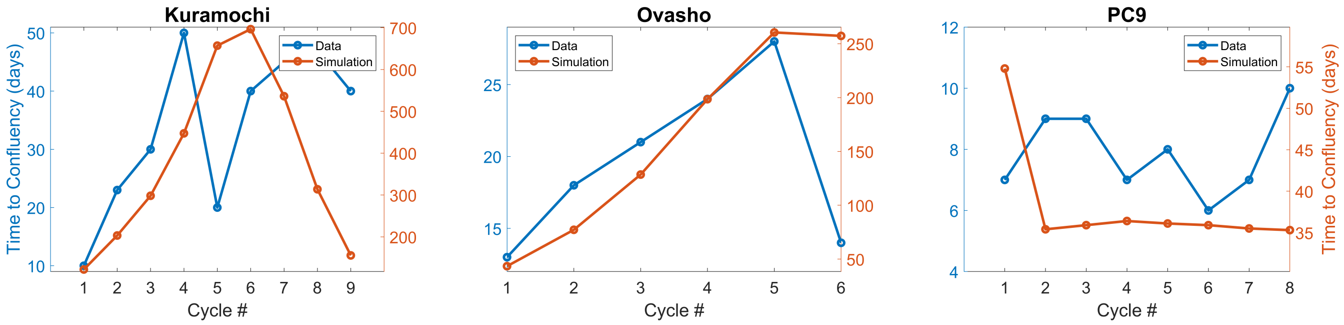


(a) (b) (c)

Figure 1: Times to confluence from França et al. [1] (blue) and calculated with our simulation for (a) Kuramochi and (b) Ovasho high-grade serous ovarian cancer and (c) PC9 non-small cell lung cancer cell lines.

Using our model, we simulated the dose-escalation experiments described in França et al. [1] for all five cell lines tested in the paper: Kuramochi, Ovasho, COV362, A375, and PC9. For each cell line, the first dose was selected (the lowest dose for the first cycle) by finding the drug concentration that most closely matches the viability (of the first dose) reported in the paper. Note that different cell lines have different dose-response protocols and the time at which the viability assay is performed. After finding the first dose, we increased the dose for each subsequent cycle by the same proportions used by the authors. We manually adjusted the parameters (Table 2) until we could qualitatively approximate a trend similar to the one observed and reported by França et al. [1] (Figure 1).

Table 2: Parameters used during simulations of the dose escalation experiment

| | $\boldsymbol{c_R}$ | $\boldsymbol{\alpha}$ | $\boldsymbol{s_r}$ | $\boldsymbol{c_v}$ |
|---|---|---|---|---|
| **Kuramochi** | $5 \times 10^{-5}$ | 0.5 | 5 | $1.25 \times 10^{-4}$ |
| **Ovasho** | $5 \times 10^{-4}$ | 1.5 | 2.5 | $1.25 \times 10^{-4}$ |
| **COV362** | $5 \times 10^{-5}$ | 0.5 | 2.5 | $9 \times 10^{-4}$ |
| **A375** | $5 \times 10^{-5}$ | 0.5 | 2.5 | $2.5 \times 10^{-3}$ |
| **PC9** | $1 \times 10^{-3}$ | 0.5 | 2 | $2.5 \times 10^{-3}$ |

### 3 Verification of Airy Distribution in $P_x(x)$

Suppose a sufficient amount of time had passed to allow $P(x, s, t)$ to reach its steady-state given by Equation 6 from the main text. Under these conditions, we focus exclusively on the $x$-dynamics described by

$$\partial_t P_x(x,t) = \mu \partial_x^2 P_x + [\xi - ax + r_x g(t)]\, P_x \tag{1}$$

where, this time,

$$g(t) = 1 - \frac{1}{N_{ss}} \int_0^1 dx\, P_x(x,t)$$

Equation 1 can be rewritten in terms of normalized probability density $p_x(x,t) := P_x(x,t) / \int_0^1 dx\, P_x(x,t)$.

$$\partial_t p_x(x,t) = \mu \partial_x^2 p_x - a\,(x - \langle x \rangle(t))\, p_x, \qquad \langle x \rangle(t) = \int_0^1 dx\, x\, p_x(x,t) \tag{2}$$

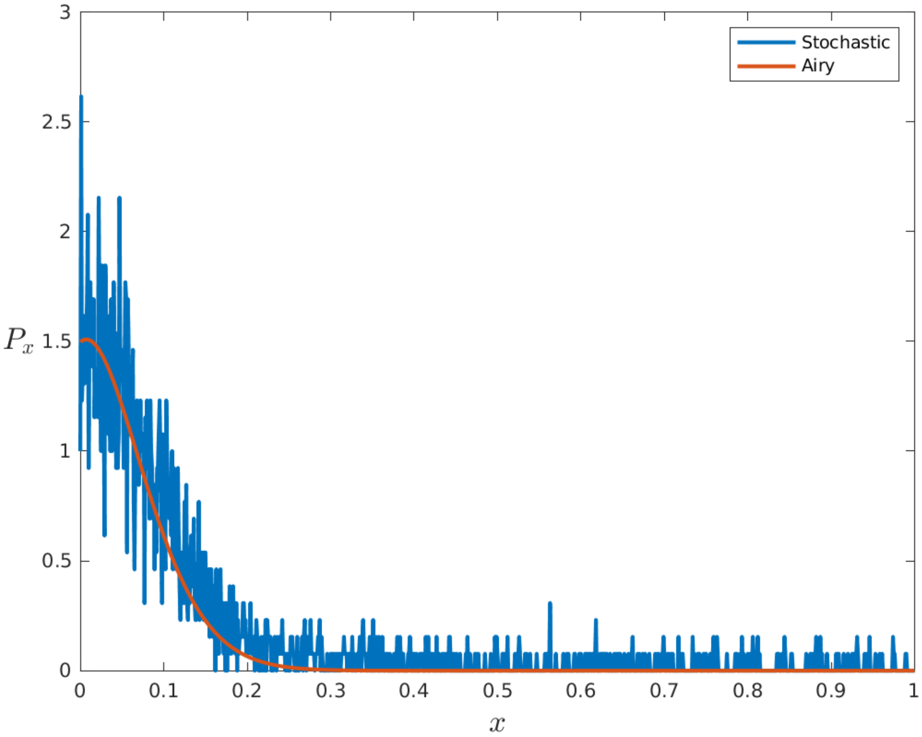


Figure 2: Stochastic simulation of the $x$-distribution compared with the analytical steady-state solution of Equation 3 from the main text.

The steady-state solution of Equation 2 is indeed an Airy distribution identical to Equation 5 from the main text. However, the Airy function is defined on all reals, whereas our operational definition of CTP confines $x$ to the interval $(0,1)$. Also, the dynamics near the boundaries, $x = 0$ and $x = 1$, on this abstract epigenetic space are ambiguous as it is biologically unclear whether the motion of a cell's state is absorbed into or reflected from these limits. Furthermore, note that $s_0 \to -\infty$ as $x \to 0$ and $s_0 \to \infty$ as $x \to 1$. Hence, to validate our assumption about the initial condition of $P(x,s,t)$ prior to treatment, we performed a stochastic simulation using the Gillespie algorithm.

The detailed scheme of the algorithm is as follows. First, the epigenetic space in $x \in (0,1)$ is discretized into units of $\Delta x$. Starting with an initial population of $N_0$, where all cells occupy the same state ($x = x_0$), the simulation is run for a sufficiently long duration $\mathcal{T}$, to allow the distribution to approach a near steady-state. During the simulation, each cell can move up or down in $x$ by $\Delta x$ at a rate of $\mu/2\Delta x^2$ while dying at a rate of $ax$. At the boundaries ($x = 0$ and $x = 1$), cells are constrained to move up or down, respectively. Because we are interested in the normalized distribution, we maintain a constant population size by placing a new cell at a random value of $x$ whenever a death event occurs. Using $N_0 = 2500$, $x_0 = 0.1$, 4000 time steps, and $\Delta x = 0.001$ (and with $\mu$ and $a$ values consistent with Table 1), we successfully replicated a sharp Airy-like distribution in $x$, closely resembling the curve generated from Equation 5 from the main text.
